# Anti-CD117 CAR T cells incorporating a safety switch eradicate human acute myeloid leukemia and hematopoietic stem cells

**DOI:** 10.1101/2023.06.15.544946

**Authors:** Chiara F. Magnani, Renier Myburgh, Silvan Brunn, Morgane Chambovey, Marianna Ponzo, Laura Volta, Christian Pellegrino, Steve Pascolo, Csaba Miskey, Zoltán Ivics, Judith A. Shizuru, Dario Neri, Markus G. Manz

## Abstract

Acute Myeloid Leukemia originates from the accumulation of mutations in hematopoietic stem and progenitor cells, leading to the emergence of leukemia-initiating cells, which sustain blast formation. CAR T cells specific for the CD117 antigen can deplete malignant and healthy hematopoietic stem cells. Here we exploit non-viral technology to achieve early termination of CAR T cell activity to prevent incoming graft rejection. Transient expression of an anti-CD117 CAR by mRNA conferred T cells the ability to eliminate CD117+ targets in vitro and in vivo. As an alternative approach, we used a Sleeping Beauty transposon vector for the generation of CAR T cells incorporating an inducible Caspase 9 safety switch. Stable CAR expression was associated with high proportion of T memory stem cells, low levels of exhaustion markers, and potent cellular cytotoxicity. Anti-CD117 CAR T cells mediated depletion of leukemic cells and healthy hematopoietic stem cells in NSG mice reconstituted with human leukemia or CD34+ cord blood cells, respectively, and could be chemically terminated in vivo. The use of a non-viral technology to control CAR T cell pharmacokinetic properties is attractive for a first-in-human study in patients with acute myeloid leukemia prior to hematopoietic stem cell transplantation.

## Introduction

Acute myeloid leukemia (AML) has a dismal prognosis due to the high rate of relapse, even after allogeneic hematopoietic stem cell transplantation (allo-SCT), which represents the therapy with highest curative potential for patients with adverse-risk disease ^1^. AML arises from the accumulation of mutations in hematopoietic stem and progenitor cells (HSPC), leading to the emergence of a population of malignant leukemia-initiating cells (LIC) ^2–4^. AML-LICs maintain high phenotypic similarity to their cells-of-origin and persisting LICs are sources of post-treatment disease relapse. Successfully targeting AML-LIC is therefore essential for AML cure. However, similar or same target expression on cell-of-origin HSPCs bares the risk and consequence of healthy HSPC depletion and subsequent severe myeloablation ^5^.

Chimeric antigen receptor (CAR) T-cell immunotherapy is a powerful treatment strategy that utilizes genetic engineering to enhance T-cell antitumor activity through ectopic expression of a receptor recognizing tumor-associated antigens ^6,7^. MHC-unrestricted recognition of antigens widely expressed on tumor cells and a multifactorial, self-sustaining T cell-mediated immune response are among the keys to the proven success in B-cell malignancies, which has led to the approval and commercialization of several CAR T products in less than five years. In all of the approved instances, the surface targets are not specific for malignant cells but are also expressed on the cell-of-origin, i.e. B- and plasma cells, which both are not key for the immediate survival of patients and which might re-grow from their upstream progenitor cells. Using a similar approach targeting cell-of-origin antigens in AML-LIC will lead to long-lasting or permanent myeloablation. Indeed, most AML antigens currently targeted by CAR T cells, such as CD123 ^8^, CD33 ^9^, and CLL-1 ^10^, are expressed by HSPCs. Therefore, they are currently being evaluated mainly in a bridge-to-transplant setting, where hematopoiesis is subsequently rescued by allo-SCT ^11–13^. This underlines the importance of either identifying specific AML-LIC targets or design strategies to mitigate expected toxicities.

We recently described the use of CAR T cells specific for the CD117 antigen to deplete LIC and replace HSPC by allo-SCT ^14^. CD117, also referred to as c-Kit, is the receptor for stem cell factor (SCF), regulating HSPC self-renewal, quiescence and survival in healthy and malignant hematopoiesis ^15–18^. Depletion of CD117+ cells may enable concomitant AML-LIC and HSPC eradication, facilitating bone marrow niche clearance and subsequent allo-SCT in absence of non-genotoxic preconditioning ^19^. This concept implies early termination of CAR T-cell activity to prevent subsequent graft rejection.

In the current study, we therefore exploit several non-viral technologies for the generation of engineered T cells temporary expressing anti-CD117 CAR molecules as well as safety-switches to pre-clinically test their efficacy and termination. Our data support clinical testing of CD117 CAR T cells in patients with relapsed AML and Myelodysplastic Syndrome (MDS).

## Results

### Transient CD117 CAR T cells generated by mRNA electroporation

mRNA incorporation into cells leads to transient protein expression. To improve the safety of anti-CD117 CAR T cells and allow for subsequent HSCT engraftment, we used mRNA electroporation to generate transient anti-CD117 CAR T cells. The protocol to deliver mRNA in T cells was established by using nucleofection technology, which we have previously used to generate transposon-based non-viral CAR T cells for B-ALL patients ^20^ and which allows highly efficient transfection (Figure 1A). Purified T cells were activated for four days and subsequently electroporated with CD117 CAR encoding in vitro transcribed (ivt) mRNA modified with pseudouridine and 5-methylcytosin (m5C). We used a previously published anti-CD117 single-chain fragment variable (scFv) ^14^ generated from the sequence of the 79D antibody ^21^. The CAR also included the hinge and transmembrane domains of the CD8 alpha chain, the 41BB costimulatory domain and the CD3ζ domain. Electroporation of T cells with mRNA CAR minimally affected the CD4/CD8 ratio and the memory phenotype of CAR T cells compared to control T cells (Figure S1). The decay kinetics of CAR expression was evaluated over time by flow cytometry on total electroporated T cells, by labeling with *Staphylococcus aureus* Protein A, which binds to antibody fragments featuring a variable heavy chain domain originating from the VH3 family^22^ (Figure 1B) or, alternatively, a target c-kit antigen (Figure 1C and 1D). High level of CAR expression was seen at 24 hours after ivt mRNA electroporation, with a mean fluorescence intensity (MFI) and a percentage of CAR-positive cells comparable to the one achieved by using a CD117 CAR encoding lentiviral vectors (LV). A progressive decrease of CAR expression was seen over time and was more evident when 1.5 μg of mRNA CAR were electroporated compared to 10 μg, showing a persisting CAR expression, thought at low level, until day 3 after electroporation (Figure 1E). We therefore selected the electroporation with 10 μg of mRNA CAR for subsequent evaluation of CAR T cell functionality. T cell viability was 73.9% (±6.2) and 87.6% (±0.8) at one and three days post electroporation, respectively (Figure 1F). Functionality of T cells electroporated with mRNA CAR was assessed at day 1, day 2, and day 3 post electroporation. Cytotoxic assays on HL-60 cell lines transduced to express CD117 in comparison to wild-type (wt) HL-60 cell line confirmed specific killing of antigen-positive cell lines by T cells electroporated with ivt mRNA CAR and absence of activity toward the CD117 negative target. Cytotoxic activity decreased over time and was correlated with CAR expression kinetics (Figure 1G). Furthermore, to evaluate CAR expression in parallel to antigen-driven proliferation, T cells electroporated with mRNA CAR were stained with 5-(and 6)-Carboxyfluorescein diacetate succinimidyl ester (CFSE) and co-cultured with target cells for 72 hours. Progressive loss of CAR expression was associated with CSFE dilution, with complete absence of CAR expression on high-proliferating CSFE^low^ cells. Complete elimination of the CD117+ target cells was observed at the end of the co-culture, indicating that transient CAR expression is sufficient for target cell eradication (Figure 1H). Having demonstrated potent in vitro activity of transient anti-CD117 CAR T cells, we evaluated their in vivo activity against CD117+ healthy human HSPCs. Transplantation of cord-blood (CB) derived purified hCD34+ cells into irradiated newborn immunodeficient mice through intrahepatic injection allows for generation of humanized mice models with lymphohematopoietic reconstitution. After establishment of human hematopoiesis in NSG mice, we treated the mice with a single dose of 2 x 10^6^ or two high doses of 6 x 10^6^ ivt mRNA CAR T cells (Figure 1I). In contrast to treatment with a single dose of CAR T cells, two subsequent high doses (3 days apart) resulted in CD117+CD33+ cell depletion in the bone marrow (BM), similarly to mice treated with CAR T cells produced by LV vectors (Figure 1L and Figure S2). In parallel with the observed target cell elimination, accumulation of CD3+ cells in the peripheral blood (PB) was observed (Figure 1J-K).

**Figure 1.**
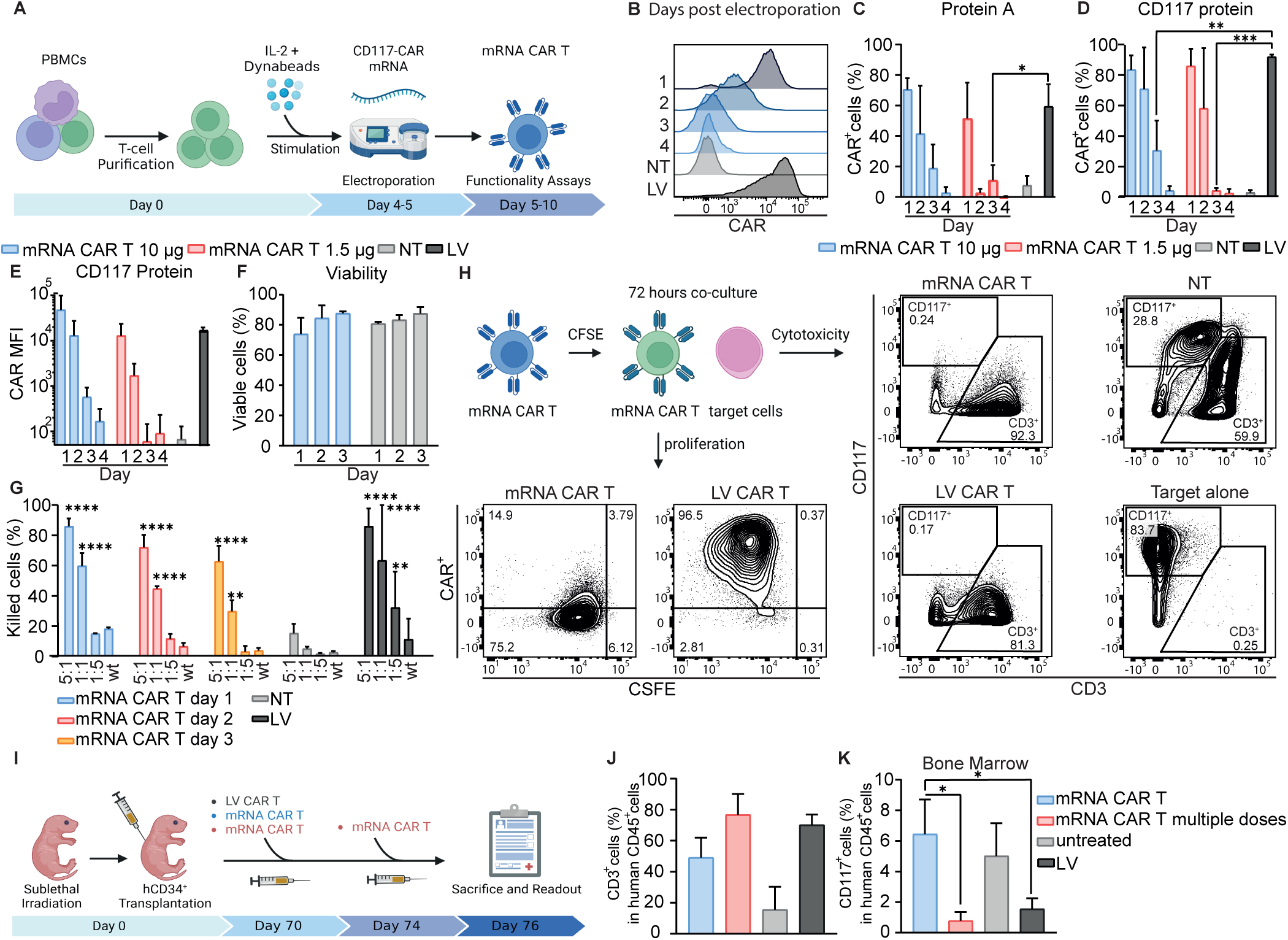
Transiently expressed CD117 CAR T cells demonstrate efficient killing in vitro and in vivo. (A) Schematic representation of the method to generate mRNA CAR T cells. (B) CAR expression of T cells electroporated with 10 µg of CAR mRNA over time. One representative histogram of results from five donors is shown. (C) Percentage of CAR+ cells at different time points after electroporation as determined by flow cytometry with Protein A. Two-way Anova with Turkey multiple comparison (D) Percentage of CD3+CAR+ cells at different time points after electroporation as determined by flow cytometry with the recombinant c-Kit protein. Two-way Anova with Turkey multiple comparison (E) CAR expression level (geometric mean) at different time points after electroporation as determined by flow cytometry with the recombinant c-Kit protein. Data illustrate the mean ±SD from 5 different donors for 10 μg, LV and NT cells, and 3 donors for 1.5 μg. (F) Viability of T cells electroporated with 10 µg of CAR mRNA over time as determined by Annexin V and 7AAD staining. Data illustrate the mean ±SD from 3 different donors. (G) Cytotoxicity of mRNA CAR T cells at different time points after electroporation compared to NT and LV CAR T cells against CD117+ HLA-60 at different E:T ratios and wt HL-60 (E:T=1:1). Data illustrate the mean ±SD from 3 different donors. Two-way Anova with Turkey multiple comparison (*compared with NT cells). (H) Schematic outline of the co-culture experiment with CFSE stained T cells. HL-60 target cells were co-culture with mRNA CAR T cells, LV CAR T cells, or NT cells (E:T 1:1) for three days. Representative Flow cytometric immunophenotyping showing elimination of target cells (cytotoxicity, gated in live cells), and proliferation of CAR T cells (gated in CD3+ cells). One representative donor of results from three donors is shown. (I) Schematic outline of the in vivo experiment. Newborn NSG mice were sub-lethally irradiated and injected with CB-derived hCD34+ cells (day 0). At day 70, mice received 2 x 106 mRNA CAR T cells, 6 x 106 mRNA CAR T cells, or 2 x 106 LV CAR T cells. Mice treated with 6 x 106 mRNA CAR T cells received an additional dose after 3 days. (J) Percentage of CD3+ cells in BM of all treated groups at endpoint. (K) Percentage of BM CD33+CD117+ cells cells of all treated group at endpoint. Data illustrate he mean ±SD from 3 different mice for mRNA CAR T cells. One-way Anova with Turkey multiple comparison.

Collectively, these data show that electroporation of T cells with anti-CD117 CAR-encoding mRNA leads to transient CAR expression and concomitant acquisition of in vitro effector functions, which are progressively lost after T cell proliferation. In addition, transient expression of the CAR molecule induces specific activity in vivo, but requires infusion of at least two high doses of CAR T cells.

### Production and characterization of non-viral CD117 CAR T cells including the iC9 switch by SB vector

Although T cells transiently expressing CD117 CAR from electroporated ivt mRNA have shown attractive features, such as efficacy and transient, self-terminating CAR expression upon proliferation, the need of at least two high doses may compromise feasibility for translation to the clinic. In order to facilitate the development of an anti-CD117 CAR strategy, we thus designed a genomically integrating vector based on a pT4 SB transposon ^23^ that includes the inducible Caspase 9 (iC9) switch and the anti-CD117 CAR, separated by a 2A peptide ^24^, under the control of the human elongation factor 1 alpha (EF-1α) promoter. SB allows the generation of CAR T cells for clinical application with demonstrated potent anti-leukemic activity ^20^. iC9 allows for rapid termination of CAR T cells by activation of the apoptotic pathway by application of a small molecule that acts as a chemical inducer of dimerization (CID) ^25^. The resulting vector has an optimized donor vector architecture, consists of an integrated cassette of 5 Kb, and allows for the stoichiometric expression of the two transgenes (Figure 2A). Our previous LV vector including the CD117 CAR and RQR8 ^26^ was used in parallel to compare SB-generated CAR T cells with CAR T cells transduced with LV. The hyperactive SB100X transposase, supplied as plasmid DNA or mRNA, catalyzes transgene integration ^27,28^.

**Figure 2:**
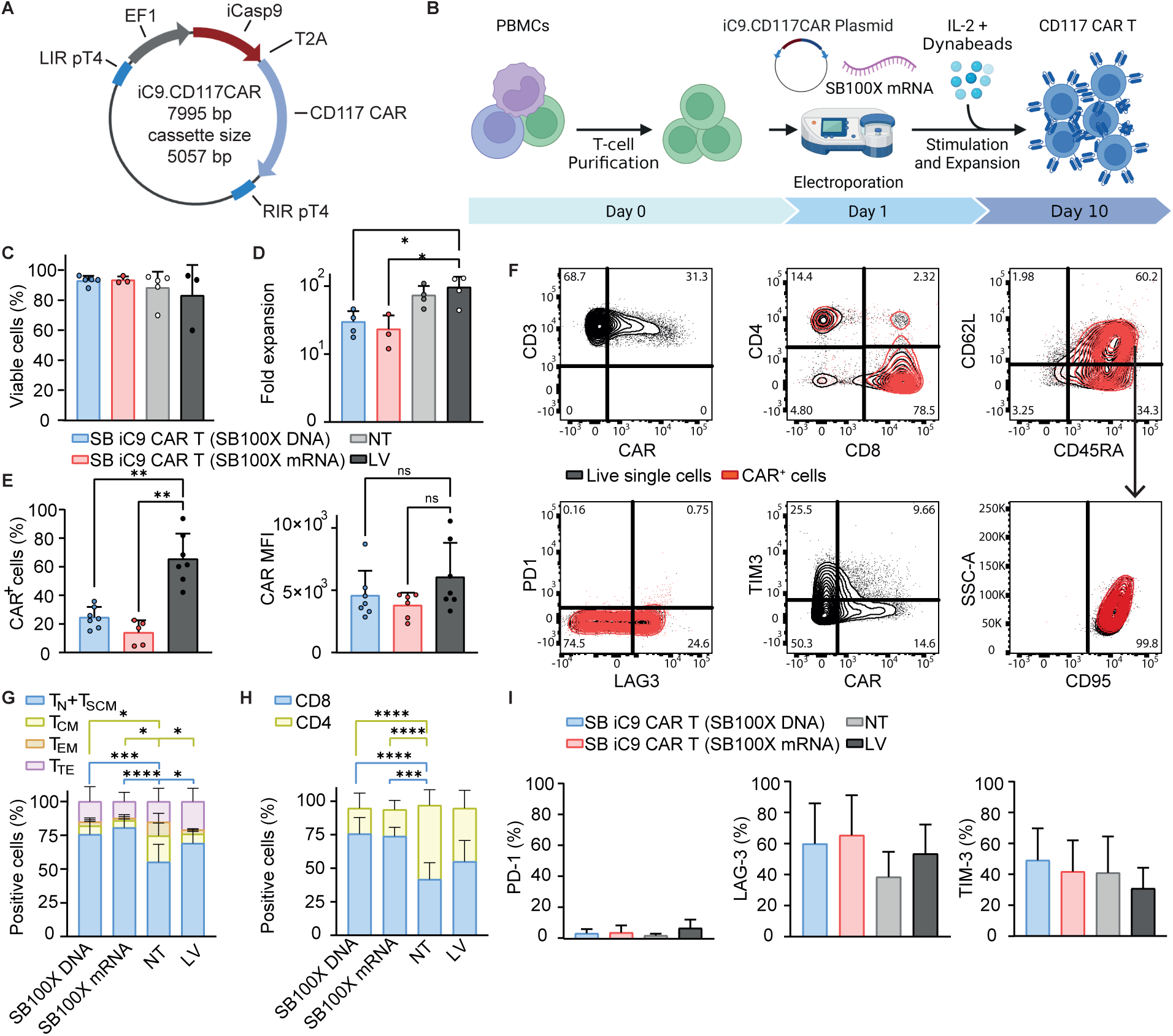
Generation and characterization of anti-CD117 CAR T cells incorporating iC9 by SB vector. (A) Schematic representation of the SB vector containing a bicistronic construct with the iC9 and CD117 CAR transgenes under the regulation of hEF-1α. The transgenes are linked by a T2A peptide. The expression cassette is enclosed in the PT4 left/right inverted repeats/directed repeats (LIR/RIR). (B) Schematic representation of the method to generate SB CAR T cells. CAR T cells were produced by electroporation with the SB vector and the SB100X transposase. (C) Cell viability at the end of the manufacturing process. (D) Fold expansion calculated by dividing the number of total viable cells at the end of the expansion phase by the respective number on day 0. (E) CAR expression as percentage of CD3 cells and expression level (geometric mean) as determined by flow cytometry with the recombinant c-Kit protein. (F) Flow cytometric immunophenotyping by dual-density plots in 1 representative batch (n = 6). (G) Percentage of CD3+CD45RA+CD62L+CD95+ naïve-like (TN) and stem cell memory (TSCM), CD3+CD45RA-CD62L+ central memory (TCM), CD3+CD45RA-CD62L-effector memory (TEM), and CD3+CD45RA+CD62L-terminal effector (TTE). (H) Percentage of CD3+CD8+ and CD3+CD4+. (I) Percentage of CD3+PD-1+, CD3+TIM-3+ and CD3+LAG-3+. Data illustrate the mean ±SD from 6 different donors. Two-way Anova with Turkey multiple comparison (*compared with NT cells).

With the purpose of transfection optimization, we compared total peripheral blood mononuclear cells (PBMC) and negatively selected bead-purified T cells as starting material. Total PBMC were stimulated with anti-CD3 antibody OKT3 as we did for producing CAR T cells for clinical application (NCT03389035). Purified T cells were stimulated with anti-CD3/CD28 paramagnetic beads and standard LV transduction was performed in parallel^14^. The protocol achieved T cell transfection with electroporation of total PBMC as well as purified T cells (Figure S3A). Since in the clinical protocol we stimulated T cells with an autologous irradiated PBMC feeder after electroporation as a source of antigen-presenting cells (APC), we evaluated whether the addition of APC can be avoided by stimulating purified T cells the day after electroporation with beads. T cell transfection was obtained in both conditions, in the presence or in the absence of PBMC as a source of APC (Figure S3B). Furthermore, we performed electroporation in the presence of different concentration of CAR transposon and SB100X plasmids or by using SB100X coding ivt mRNA. We also tested a condition with electroporation performed two days after stimulation. In our hands, electroporation of purified T cells with 7.5 ug of CAR transposon vector and 2.5 ug of SB100X DNA, followed by bead stimulation allowed for the highest transduction efficiency (Figure S3C) and a favoured in vitro expansion calculated as population doubling (Figure S3D) at the end of a 10-days in vitro culture (Figure 2B). SB100X could also be provided with ivt mRNA, particularly using the highest tested dose of 15 ug for 5X10^6^ total T cells.

SB iC9.CAR T cells generated with the optimized conditions had a high level of viability (mean 93.1% ± SD 3.0%; range, 88.0%-96.4% for CAR T cells generated with SB100X DNA, and mean 93.5% ± SD 2.3%; range, 92%-96.2% for CAR T cells generated with SB100X mRNA) (Figure 2C), and a mean fold T cell number increase of 29.9 (range, 17.6-46.0), supporting the production of clinically relevant numbers of non-viral CAR T cells (Figure 2D). CAR T cells showed stable CAR expression which was characterized by a mean fluorescence intensity similar to the one achieved after LV-mediated transfection (Figure 2E). To understand whether gene transfer methods affect the final T cell composition, we performed extensive phenotyping at the end of differentiation. Notably, the procedure of generating CAR T cells with SB transposon did not affect the memory differentiation of T cells (Figure 2F). SB iC9.CAR T cells retained a higher proportion of CD45RA+CD62L+ naïve-like population with features of T stem cell memory (mean 75.6% ± SD 9.6%, and mean 80.7% ± SD 6.4% for CAR T cells generated with SB100X DNA and SB100X mRNA, respectively), counterbalanced by a lower percentage of CD45RA-CD62L+ central memory (mean 6.4% ± SD 5.6%, and mean 5.3% ± SD 4.2% for CAR T generated with SB100X DNA and SB100X mRNA, respectively) in comparison to non-transduced (NT) cells (Figure 2G). There was an increase in CD8/CD4 proportion in SB-generated CAR T cells compared CAR T cells transduced with LV (mean of CD8+ cells 75.6% ± SD 12.0% and 73.8% ± SD 6.7% for CAR T cells generated with SB100X DNA and mRNA, respectively, compared to 64.9% ± SD 15.7% in LV-transduced CAR T cells, Figure 2H). Moreover, CAR T cells generated with SB transposon showed low levels of the exhaustion markers PD-1 (mean 3.0%, SD 2.8%), LAG3 (mean 59.9%, SD 26.1%), and TIM3 (mean 49.2%, SD 20.6, Figure 2I). In sum, these results demonstrate the potential of non-viral technology to efficiently deliver large multi-cistronic vectors and allow for simultaneous expression of multiple transgenes without affecting the viability and quality of primary T cells, supporting large-scale production and utilization at clinical level.

### SB iC9.CAR T cells exhibit antigen specific effector functions and iC9-mediated elimination of gene-modified cells

We subsequently tested whether generation of CAR T cells by non-viral engineering of a large, bi-cistronic cassette impacts on the functional activity in vitro. To evaluate their cytotoxic activity, SB iC9.CAR T cells were co-cultured for 24 hours with the AML cell line MOLM-14, which was transduced, and sorted to express human CD117, luciferase, and GFP. SB iC9.CAR T cells exhibited potent cytotoxic activity against the target, similarly to cells engineered with LV encoding for the same CAR. Potent cytotoxicity was observed with unpurified CAR T cells at an Effector:Target (E:T) ratio of 1:1 (Figure 3A) and with T cells sorted for CAR expression, even in conditions where CAR T cells are at a disadvantage with respect to the target cells at E:T ratio of 1:5 (Figure 3B). Because the inferior amount of CAR+ T cells generated with SB compared with cells transduced with LV may lead to decreased functionality, we analysed the level of pro-inflammatory cytokine production after target recognition in mixed CAR+ and CAR-T cells not purified for CAR expression. CAR T cells generated with SB or LV vectors did not differ in their ability to secrete interferon (IFN)-γ, tumor necrosis factor (TNF)-α, and interleukin (IL)-2 (Figure 3C). Having proved specific and comparable effector activity of CAR T cells, we then evaluated if they are rapidly terminated by activation of the inducible Caspase 9. The addition of 200nM of the CID AP20187 to cultures of anti-CD117 CAR T cells induced apoptosis of > 95% of transduced CAR T cells within 24h, but had no effect on the viability of NT cells (Figure 3D). This was also true with CAR+ sorted cells and even at low amount of the CID (Figure 3E). Analysis of the transgene copy number of CAR T cells generated by SB revealed a mean of 2.5 (range 1.9-3) transgene integrations per cell (Figure S4). We conclude that SB iC9.CAR T cells are cytotoxic against the MOLM-14 cell line and are able to release cytokines. Delivery of iC9 transgene by non-viral technology is feasible and functional to equip CAR T cells with a suicide gene as a safety measure.

**Figure 3:**
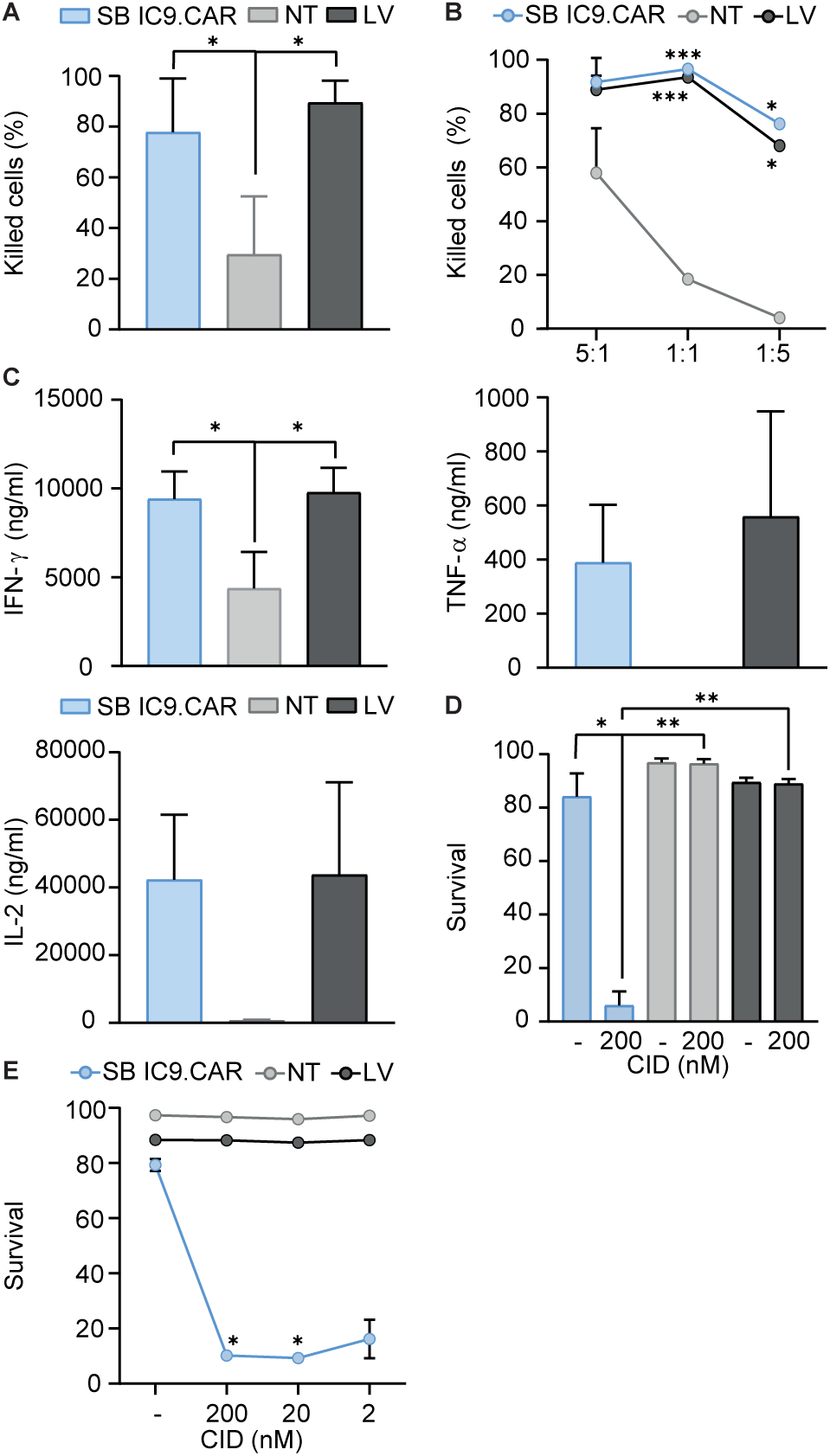
anti-CD117.iC9 CAR-T cells exhibit potent functionality and are rapidly terminated by iC9 activation. (A) Killing activity of CAR T cells after 24 hours of co-culture with CD117+ Molm-14 at an E:T ratio of 1:1. Data illustrate the mean ±SD from 3 different donors. One-way Anova with Turkey multiple comparison (*compared with NT cells). (B) Killing activity of sorted CAR T cells at different E:T ratios. One representative experiment of results from three donors is shown. (C) IFN-γ, TNF-α, and IL-2 secretion of CAR T cells after 24 hours of co-culture with CD117+ Molm-14. Data illustrate the mean ±SD from 3 different donors. One-way Anova with Turkey multiple comparison (*compared with NT cells). (D) Apoptosis assessed in CAR T cells in the presence of 200 nM CID. Data illustrate the mean ±SD from 3 different donors. One-way Anova with Turkey multiple comparison (*compared with NT cells). (E) Apoptosis assessed in sorted CAR T cells at different CID concentrations. One representative experiment of results from three donors is shown.

### SB iC9.CAR T cells show potent anti-leukemic activity in vivo

To evaluate the activity of iC9.CAR T cells in an AML mouse model, sub-lethally irradiated immunodeficient NOD.Cg-Prkdcscid Il2rgtm1Wjl/SzJ (NSG) mice were transplanted with 0.1X10^6^ MOLM-14 cells modified to express high level of CD117 and luciferase. CAR T cells were injected intravenously at day 9 after MOLM-14 infusion, when all the mice had measurable sign of tumor engraftment (Figure 4A). CAR T cells are generally infused as bulk population for clinical application. Therefore, to evaluate the same dose of CAR T cells, we decided to infuse 2X10^6^ CAR+ T cells per mouse, corresponding to 7.4X10^6^ SB iC9.CAR T cells and 3.3X10^6^ LV CAR T cells. As controls, mice were infused with PBS or treated with 7.4X10^6^ NT cells. SB iC9.CAR T cells mediated robust anti-leukemic activity in vivo, as demonstrated by the flux signal, leading to a significant survival advantage over control mice (median overall survival for NT= 22.5 days vs. SB= not reached, p= 0.0091, Mantel-Cox) (Figure 4B-D). Notably, SB iC9 CAR T cells were as efficient as CAR T cells transduced with LV (median overall survival for NT= 22.5 days vs. LV= not reached, p= 0.0091, Mantel-Cox). At the time of sacrifice, mice treated with SB iC9.CAR T cells showed almost complete clearance of leukemic blasts in the BM (Figure 3E) and spleen (Figure 3F). In this model, we observed a residual flux signal associated with circulating tumor blasts in the PB of 4 out of 4 mice treated with LV CAR and 2 out of 4 mice treated with SB iC9 CAR. Taken together, these data indicate a comparably strong anti-leukemia activity in a xenograft model of human AML with LV and SB anti-CD117 CAR T cells.

**Figure 4:**
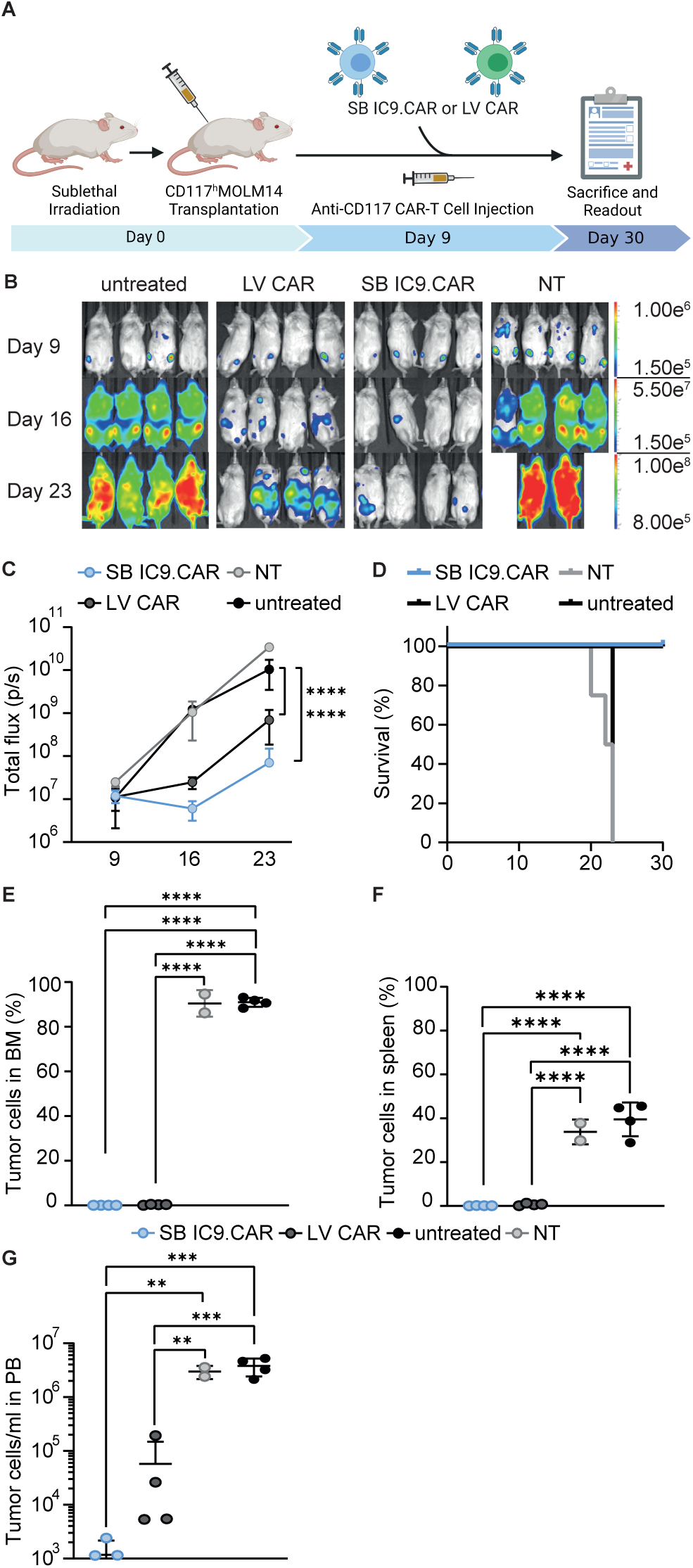
SB-generated anti-CD117 CAR-T cells show potent anti-leukemic activity in vivo. (A) Schematic representation of the in vivo experiment. NSG mice were sub-lethally irradiated and injected with MOLM-14 CD117+ Luciferase+ cells. At day 9, mice were injected with 2X106 CAR T cells, generated with SB or LV. (B) IVIS images of mice injected with CAR T cells, NT cells or left untreated. (C) Bioluminescence intensity (total flux per second) over time in mice receiving the indicated treatment. Two-way Anova with Turkey multiple comparison (*compared with untreated). (D) Kaplan-Meier survival curves over time. Statistical differences are calculated by Mantel-Cox test. (E) Flow cytometry analyses of BM at sacrifice, corresponding to day 20-23 for control groups (humane endpoints), and day 30 for groups treated with CAR T cells. (F) Flow cytometry analyses of spleen at sacrifice. (G) Flow cytometry analyses of PB at sacrifice. Data illustrate the mean ±SD from 4 different mice per group. One-way Anova with Turkey multiple comparison.

### Elimination of SB iC9.CAR T cells in vivo after HSPC depletion to enable subsequent allo-SCT

CD117 targeting is attracting increasing interest to eliminate healthy human HSPCs as a non-cytotoxic conditioning regimen before transplantation ^19,29^. To examine the ability of SB iC9.CAR T cells to deplete healthy HSPC in vivo, irradiated newborn immunodeficient mice were reconstituted with human CD34+ cord blood cells (Figure S5A). On day 40 after transplantation, mice showed a mean of 27.6% hCD45+ PB cell engraftment, but absence of mature human cells in PB (Figure S5A, B). Mice then received 2X10^6^ SB iC9 CAR T cells and were analyzed after two weeks of treatment. SB iC9.CAR T cells decreased CD117+ cell engraftment in the BM (1.1%±0.5) compared with untreated animals, which showed a mean of 5.6% (±0.8) of CD117+ HSPCs (Figure 5B). An increase of CD3+ cells in the BM of treated animals (Figure 5B) was associated with the observed reduction of CD117+ cells (Figure 5C-D).

**Figure 5:**
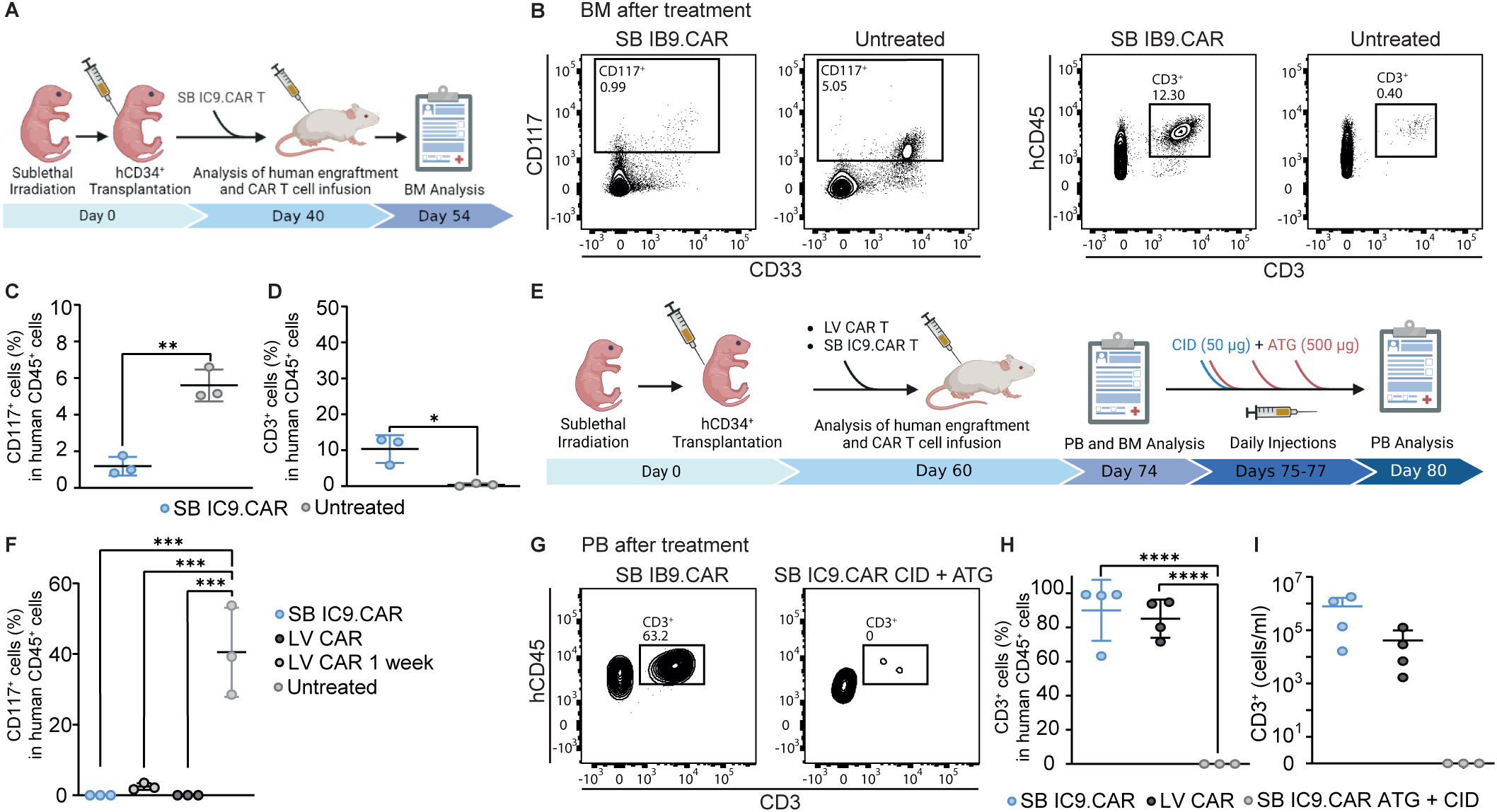
SB-generated CAR T cells are able to deplete human CD117+ HSPC and are terminated by iC9 activation and ATG. (A) Newborn mice were sub-lethally irradiated and injected intrahepatically with human CB-derived CD34+ cells. Following engraftment at day 40, mice were injected with 2X106 CAR T cells generated with SB. (B) Representative flow cytometry plots of CD117+ and CD3+ cells in BM of animals from each treatment group after two weeks of treatment. (C) Percentage of CD117+ of hCD45+ cells in BM. (D) Percentage of CD3+ cells of hCD45+ cells in BM. (E) Newborn mice were sub-lethally irradiated and injected intrahepatically with human CB-derived CD34+ cells. Following engraftment at day 60, mice were injected with 2X106 CAR T cells generated with SB or LV. Mice were then treated with a single dose of CID (50ug/mouse) and three subsequent doses of ATG (500ug/mouse) from day 75 to day 77. (F) Percentage of CD117+ cells of hCD45+ cells in BM after one or two weeks of treatment. (G) Representative flow cytometry plots of hCD3+ cells in PB of animals from each treatment group before and after CAR T termination. (H) Percentage of CD3+ cells of hCD45+ cells and (I) CD3+ cells/ml in PB before and after CAR T termination.

Complete elimination of CD117+ HSPCs would allow clearance of malignant LIC and healthy HSPCs, but the persistence of CD117-targeting CAR T cells would compromise subsequent allo-SCT. Having demonstrated that SB iC9.CAR T cells efficiently target healthy CD117+ cells in humanized mice, we assessed the possibility to terminate CAR T cells after the complete elimination of CD117+ HSPCs. We consequently generated humanized mice by engrafting human HSPCs and waited 60 days for high human cell engraftment (mean 27.19% ± SD 17.8) and reconstitution of mature human hematopoiesis (Figure S5C-D). In this setting, we evaluated CD117+ cell depletion after 7 and 14 days of treatment and compared once more the activity of non-viral CAR T cells with LV-engineered CAR T cells (Figure 5E). CD117 CAR T cells were able to eliminate CD117+ HSPCs after two weeks of treatment (40.5%±12.64 and 0%±0 as mean engraftment in the control group compared to treated animals), while one week of treatment reduced the engraftment (2.46±0.94) but was not sufficient to achieve complete CD117+ cell depletion (Figure 5F). High CD3+ engraftment was observed in the BM of treated animals (Figure 5G). To eliminate SB iC9.CAR T cells in vivo, mice were treated with a single dose of CID on day 17 after CAR T-cell treatment. Since mice were infused with non-sorted CAR T cells, three doses of Antithymocyte Globulin (ATG) were provided on days 15, 16, and 17 to also eliminate the fraction of not-transduced and thus non-iC9 expressing T cells, as it is usually included in allo-SCT conditioning regimen and was previously used by us to deplete CAR T cells in vivo ^14^. Treatment with the combination of CID and ATG completely eliminated anti-CD117 CAR T cells and T cells (Figure 5H-I) in the PB of respective animals at day 5 after CAR T cell termination treatment. Overall, these results demonstrate the feasibility of completely eliminating CD117 CAR T cells having mediated the healthy HSPC depletion, supporting their potential use as a conditioning regimen in allo-SCT. The analysis did not show any significant difference in the CD117+ HSPC depletion between the groups treated with LV-engineered CAR T cells and CAR T cells generated with SB.

## Discussion

We aimed to develop a CAR T cell approach to target AML-LIC that could also be used as a conditioning regimen prior to allo-SCT. We showed that electroporation of an IVT mRNA encoding a CD117 CAR is feasible to generate functional T cells that temporarily express CD117 CAR molecules, exhibit transient cytotoxicity, and in vivo activity when infused in at least two high doses. Alternatively, stable expression of CD117 CAR and the iC9 suicide gene in T cells by using a genomically integrating SB vector led to CAR expression and in vitro CAR T cell activity towards CD117+ AML cells. Activation of the iC9 transgenes induced CAR T cell apoptosis, which would allow for CAR T cell depletion before allo-SCT. Anti-CD117 CAR T cells engineered with the SB vector showed anti-leukemic activity in a human tumor xenograft model and completely depleted healthy HSPC in immunodeficient mice reconstituted with a human hematopoiesis. We observed no difference in activity when CAR T cells produced by a non-viral approach with the SB vector were compared with cells engineered with LV vectors. SB iC9.CAR T cells were terminated in vivo by combining activation of iC9 and ATG administration. AML LIC derive from HSPCs with little immune-phenotypic distinction ^2,3,30^. In contrast to mature lineage antigen-targeting in B-lineage and plasma cell neoplasia, there is no consensus on which surface antigen might be a suitable target to develop anti-AML therapies, eradicating LIC. We and others identified CD117 as potential target as it is expressed by the CD34+CD38- and CD34+CD38+ cell fractions in healthy BM and CB, and in about 80% of AML blasts ^19^. In contrast to myeloid markers as CD33, which are not required for cell survival ^31^, CD117 might be of relevance for SCF mediated LIC support ^32^, and CD117 might thus be less prone to immune-pressure selected antigen loss. As antigens suitable for targeting the AML-LIC are commonly shared by healthy HSPCs and myeloid progenitors, we evaluated whether CAR T cells targeting CD117 would eliminate HSPCs in vivo. Flow cytometric analysis of CD117+ cells in BM of animals engrafted with human hematopoiesis revealed HSPCs reduction after one week of infusion and complete eradication after two weeks. Likewise, the majority of anti-AML CAR T cell strategies have the concern to induce severe bone marrow aplasia ^33^. Preliminary clinical evaluation of CD33 CAR T cells in patients with AML indeed showed on-target/off-tumor toxicity to HSPCs and myeloid progenitors, observed as pancytopenia after treatment ^34^. Even though CLL-1 is expressed at lower levels by healthy HSPCs and has a higher specificity for AML-LIC, reduction of monocyte and neutrophil counts were observed in pediatric and adult patients treated with anti-CLL-1 CAR T cells, respectively ^35,36^. Similar or even more severe BM aplasia is expected after CD117 targeting, thus making it necessary to replace hematopoiesis with allo-SCT after immunotherapy. To spare normal HSPCs and avoid allo-SCT, target antigens recognizing AML-LIC without HSPCs toxicity such as CD70 have been proposed^37^. Nevertheless, CD70 antigen density on AML cells is below that needed to trigger CAR T cell response, which can only be achieved by increasing CAR binding avidity and pharmacological intervention^38^.

The use of an anti-AML CAR strategy as a bridge to SCT approach implies the need to mitigate toxicities on the incoming graft, i.e. graft rejection. We previously demonstrated significant in vivo depletion of CD117 CAR T cells with Rituximab using a vector incorporating the selection marker RQR8^14^, which combines target epitopes from CD34 and CD20 antigens ^26^. However, depletion of CAR T cells was incomplete. To completely eliminate CAR expression, we tested the use of ivt mRNA to manufacture transiently active CAR T cells, obviating the need to administer agents for elimination. Of note, mRNA technology has taken a huge step forward in recent years due to the Covid-19 mRNA vaccines, increasing the availability of platforms to generate mRNA for clinical application. In this study, we took advantage of recent progresses in mRNA design ^39–41^ and used modified RNA (pseudouridine and 5-methyl cytosine) as well as CleanCap®^42^ to improve translational capacity and diminish immunogenicity ^43^. The optimized mRNA design in combination with an improved protocol to generate mRNA CAR T cells by electroporating T cells after stimulation resulted in heterogeneous >85% CAR expression, with minimal loss of viability. Compared with other protocols that manipulate resting T cells to avoid early loss of CAR expression ^44^, our manufacturing protocol was faster and achieved similar kinetics in the acquisition of CAR-redirected effectors. Reducing the manipulation of CAR T cells ex vivo is known to preserve a less differentiated memory phenotype and effector activity, and indeed, most mRNA CD117 CAR T cells showed a central or effector memory phenotype. Furthermore, mRNA CD117 CAR T cells showed potent effector activity and eliminated resident HSPCs in a humanized mice model engrafted with human CD34+ CB-derived HSC. However, an important limitation emerging from our in vivo preclinical results is the need of manufacturing multiple doses, which can be challenging as observed in clinical trials with mRNA CAR T cells^45^. This limitation might be overcome through in vivo delivery of mRNA with lipid nanoparticles ^46^. However, this strategy may not be ideal for addressing a rapidly progressive disease such as AML, as suggested by the lack of demonstrated efficacy of the phase I study with CAR CD123 mRNA T cells (NCT02623582)^47^.

Based on these results, we combined stable transduced CAR T cells with the iC9 transgene, which guarantees rapid elimination of CAR T cells upon administration of the inert and approved drug Rimiducin. Moreover, selective modulation of CAR T cell expansion can be achieved by titrating the dose of the CID, which allowed mitigation of toxicities such as immune effector cell-associated neurotoxicity syndrome (ICANS) after CD19 CAR T cell treatment without interference on CAR T cell efficacy ^25^. CD117 is also expressed by mature mast cells ^48^, interstitial cells of Cajal, melanocytes, tubular epithelial cells of the kidney, and certain cells in the reproductive organs ^14^. Consequently, the presence of a switch that can induce apoptosis within minutes of administration and with a clinical proof of efficacy ^49^ might be beneficial to ensure safety and successful translation of this approach to the clinic. We demonstrated that the inclusion of iC9 in our CD117 CAR vector allows for efficient apoptosis induction in CD117 CAR T cells upon CID administration, without affecting CAR T cell effector functions evaluated in vitro and in murine xenograft models in vivo. Interestingly, mice show comparable expression of CD117 in organs and tissues^14^. The absence of reported on-target non-hematopoietic toxicity in a murine CD117 CAR in syngeneic mouse model ^50^ gives confidence that future safety evaluation will be feasible in the context of early phase clinical trials in humans.

An important observation that emerges from the data comparison is the feasibility of SB vector to allow for T cell engineering with complex and large gene cassettes. No significant differences in anti-tumor activity were identified between LV-transduced and SB-engineered CD117 CAR T cells in vitro and in vivo. We only observed a peculiar increase in CD8/CD4 proportion in SB iC9.CAR T cells, despite using the same CAR design in terms of stimulatory domains and cytokines to expand CAR T cells, which may be due to SB itself or the electroporation procedure. As previously shown for CAR T cells engineered with SB and *piggyBac* transposons ^51^, our SB iC9 CAR T cells maintain a good proportion of T stem cell memory phenotype, which is associated with superior anti-tumor activity in mouse models ^52^. Moreover, SB iC9 CAR T cells showed low levels of inhibitory receptors, which are known to contribute to T cell exhaustion and have been recently identified as predictors of cellular products associated with decreased response ^53^. We previously demonstrated the first proof of concept for the clinical safety and anti-leukemic efficacy of SB-engineered CAR T cells ^20^. Now, taken together, these results suggest that our SB multi-cistronic vector can generate CAR T cell products with a T cell composition associated with T cell fitness and potent anti-tumor activity in xenograft models. Since the CAR T cell landscape is rapidly evolving towards an increased complexity in transgene design to allow for multi-targeting, our observation clearly positions SB as valuable non-viral vector for future applications.

An increasing number of pre-clinical studies in immunocompromised mice demonstrated the utility of CD117 depletion as immunologic conditioning, allowing for HSC niche clearance and engraftment of donor-derived HSCs ^54^. CD117 antibody-mediated immunological conditioning has the advantage to avoid the toxicity of cytotoxic regimens applied in allo-HSCT, is currently being clinically evaluated in benign hematologic diseases and has shown encouraging preliminary data in terms of safety and activity ^55^. However, complete eradication of host HSCs is likely not achieved with anti-human CD117 antibodies ^29^, indicating that increased potency may be needed for hematological malignancies to allow for eradication of host HSC/LIC. We proved that SB iC9 CAR T cells are able to completely eradicate healthy HSCs in mice reconstituted with a human hematopoiesis, allowing for niche clearance and thus possibly obviating the use of cytotoxic conditioning before transplantation in AML or MDS. Notably, our data indicate that CAR T cells have to persist two weeks for achieving complete HSC/LIC depletion, which may also lead to durable leukemia remission. Patients can usually tolerate myeloablation and cytopenia expected from HSC killing for up to two weeks, indicating a possible time window for intervention. Unfortunately, our humanized model precludes the study of subsequent donor HSC engraftment because of competition with mouse HSPCs and possible rejection/GVHD by mature human T cells from the previous graft^56^. For the clinical application, we envision co-administration of CID and lymphodepleting regimen, such as ATG, which is commonly used in the clinic before transplantation, but might interfere with the SCID-repopulating ability of human CD34 cells in mouse xenografts ^57^. Using CID and ATG, we proved effective SB iC9.CAR T cell termination and complete T cell depletion in vivo.

Taken together, our results indicate that non-viral engineering of CD117 CAR T cells can lead to potent anti-tumor activity and complete eradication of HSPCs, envisioning its application prior to allo-HSCT in early clinical trials of patients with high-risk AML or MDS.

## Material and methods

### Generation of CAR mRNA

The generation of the CD117 CAR from the 79D antibody ^21^, including the CD8 hinge, CD8 transmembrane, 41BB co-stimulatory, and CD3zeta signaling domains was accomplished as previously described by us ^14^. A PCR template encoding the mRNA CD117 CAR sequence under the control of a modified T7 promoter sequence was transcribed in vitro at the mRNA platform of the University Hospital of Zürich according to a previous report ^58^. Capping and nucleotide modifications were introduced co-transcriptionally by having in the transcription reaction the CleanCap®, Pseudouridine triphosphate and eventually (for the mRNA coding the CD117 CAR) 5-Methylcytidine triphosphate. When needed the mRNA was polyadenylated after transcription using a recombinant poly-A polymerase as recommended by the manufacturer (NEB Biolabs).

### Production of the LV encoding the CD117 CAR

The viral vector encoding the anti-CD117 CAR and the RQR8 sequences previously cloned in the pCDH-EF1α-MCS-T2A-GFP lentiviral plasmid (System Biosciences, Palo Alto, CA, USA) as produced as previously reported ^14^. The RQR8 gene sequence was generously shared by Dr. Martin Pule (University College London, UK) ^26^. Briefly, HEK293T Lenti-X^TM^ cells (Takara Bio Europe, Saint-Germain-en-Laye, France) were transfected with the CAR-encoding transfer vector, psPAX2 packaging, and pCAG-VSVG envelope plasmids (both kindly provided by Dr. Patrick Salmon, University of Geneva, Switzerland) using JetPRIME® transfection reagent (Polyplus transfection, Illkirch, France). Viral particles were harvested 2 days later and concentrated with Peg-itTM (System Biosciences, Palo Alto, CA, USA).

### Generation of SB iC9.CAR vector

The pT4.iC9.79D vector, which encodes the iC9 and the CD117 CAR sequences, was obtained from the pTMNDU3 plasmid ^59^ by replacing the CD19CAR and the inverted terminal repeats (ITRs). ITRs of the pT4 vector (Addgene, 117046)^23^ and the iC9 gene ^60^ were synthesized by GeneArt (Life Technologies). The iC9 was inserted by restriction digestion into the LV under the control of a human EF1 promoter and in front of the CD117 CAR, separated by T2A element ^24^. Functional analyses were performed on bulk cell products if not otherwise specified.

### CAR T cell production with LV

CD117 CAR T cells were produced by transduction of purified T cells from healthy donors as previously described ^14^. Briefly, excess buffy coats of anonymized healthy blood donors were acquired from the Zurich blood donation service (Blutspende Zurich, Switzerland). Peripheral blood mononuclear cells (PBMCs) were isolated by density gradient centrifugation (Ficoll-Paque Plus, GE Healthcare, Chicago, Il, USA). T cells were negatively purified with EasySepTM beads (human T-cell isolation kit, STEMCELL Technologies, Vancouver, Canada). T cells were cultured in Advanced RPMI 1640 enriched with FBS (10%), Glutamax^TM^, and Penicillin/Streptomycin (100 U/ml/100µg/ml) (Gibco^®^, Thermo Fisher Scientific, Waltham, MA, USA) and stimulated with 80 U/ml IL-2 (Peprotech^®^, Rocky hill, NJ, USA). T cell activation was performed with CD3/CD28 Dynabeads^®^ (Thermo Fisher Scientific) followed by transduction with LV at MOI 3 in the presence of 8 µg/ml Hexadimethrine bromide (Sigma-Aldrich, St. Louis, MO, USA) the following day. On day 3, beads were removed with magnets. LV CAR T cells were cultured for 10 days before cryopreservation or immediate use.

### CAR T cell production with mRNA

CAR T cells were produced by electroporation of purified T cells from healthy donors (Blutspende Zurich, Switzerland). T cell activation was performed with CD3/CD28 Dynabeads^®^ (Thermo Fisher Scientific) and 80 U/ml IL-2 (Peprotech^®^, Rocky hill, NJ, USA). mRNA CD117 CAR were electroporated in T cells after 4 days post activation. Briefly, 5X10^6^ T cells were resuspended in Human T Cell Nucleofector™ Solution (Lonza) and electroporated in the presence of the CD117 CAR mRNA using the Amaxa program T-20.

### CAR T cell production with SB

SB CAR T cells were produced by adapting the previously reported method to conventional T cells ^61^. Human T cells were purified with EasySepTM beads and resuspended in Human T Cell Nucleofector™ Solution (Lonza). For each cuvette, 5X10^6^ purified T cells were electroporated (program U-14) in presence of the pT4.iC9.79D vector and the transposase SB100X plasmid (Addgene 34879) or of the transposase SB100X mRNA. After electroporation, T cells were activated with CD3/CD28 Dynabeads^®^ (Thermo Fisher Scientific) and stimulated with 80 U/ml IL-2 (Peprotech^®^, Rocky hill, NJ, USA). CAR T cells were cultured for 10 days before cryopreservation or immediate use.

### Cell lines

The HL60 and MOLM-14 cell lines were purchased from ATCC. The cell lines were cultured in RPMI 1640 supplemented with FBS (10%), and Penicillin/Streptomycin (100 U/ml/100µg/ml) (Gibco^®^, Thermo Fisher Scientific, Waltham, MA, USA). The HL-60 and MOLM-14 AML cell lines were modified to express a truncated version of the GNNK+ isoform of the *KIT* gene by LV gene transfer. The MOLM-14 AML cell line was co-transduced with the pCDH-EF1-Luciferase-T2A-GFP LV vector ^14^.

### Flow cytometry and cell sorting

Cell suspension from in vitro culture or isolated from PB, BM, or spleen were stained with antibodies in 100 μl of phosphate-buffered saline supplemented with 2% FBS for 10 minutes at room temperature. Details on the antibodies used for immunophenotyping are reported in Supplemental Table 1. Surface expression of the CD117 CAR was detected with the Recombinant human c-Kit Protein (His Tag, SinoBiological), followed by the anti-CD117 antibody. In all analyses, the population was gated based on singlet gating followed by forward and side-scatter characteristics. All subsequent gates were set according to unstained controls. Live cells were gated using Zombie Aqua™ Fixable Viability Kit (Biolegend). For cell number quantification, Flow-Count Fluorospheres were used according to the manufactory’s instructions (Beckman Coulter, Pasadena, CA). Cells were acquired with an LSR II Fortessa cytometer (BD Bioscience, Franklin Lakes, NJ, USA) and the data were analyzed with FlowJo (version 10.0.8, LLC, Ashland, USA). To purify CAR+ T cells, CAR T cells were stained with CD3 and the Recombinant human c-Kit Protein, followed by anti-CD117 antibody and sorted with a FACS Aria III (BD Biosciences).

### Cytotoxic assay

Cytotoxicity was evaluated with a 24-hour co-culture assay as previously reported. Cell death and apoptosis were detected using the GFP-Certified™ Apoptosis/Necrosis detection kit (Enzo Life Sciences, Farmingdale, NY), according to the manufactory’s instructions. T cells were co-cultured with targets at indicated E:T ratio. Target cells were identified according to GFP or were previously labeled with CFSE (1 μM, eBioscience). The final percentage of killed cells was determined as follows: (% of early necrotic and apoptotic co-cultured cells - % of early necrotic and apoptotic target cells alone) X100 / (100 - % of early necrotic and apoptotic target cells alone). All samples were run in duplicate. For long-term cytotoxic and proliferation assay, mRNA CAR T cells were stained with CellTrace™ CFSE Cell Proliferation Kit (Thermo Fisher Scientific) and co-cultured with target cells (E:T 1:1) as described previously ^62^. After three days of incubation, cells were stained for expression of CAR, CD3, and CD117 (Supplemental Table 1), and with Zombie Aqua Live/Dead (BioLegend, San Diego, CA, USA). Antibody staining followed by flow cytometry was used to evaluate target cell elimination and CD117CAR expression, while CAR T cell proliferation was determined by assessing CFSE staining dilution.

### CID-mediated induction of apoptosis

T cells were cultured in the presence of AP20187 (2-200 nM). After 24 hours, cells were stained for expression of CAR, CD3, and with the Apoptosis/Necrosis detection kit (Enzo Life Sciences). Apoptosis induction was assessed with flow cytometry. Survival was calculated by analyzing the percentage of viable residual cells in CD3+CAR+ cells considering the proportion of CAR+ cells,in the presence of AP20187 versus medium alone as follows: (% of CAR+ alive cells) * (1 – (CAR+ cells with medium – CAR+ cells with AP20187)/(CAR+ cells with medium)).

### Cytokine release assays

T cells/ml were stimulated with leukemic blasts at a ratio of 1:1. After 24 h, culture supernatants were harvested and levels of INF-γ, TNF-α, and IL-2 were investigated with an enzyme-linked immunosorbent assay (ELISA MAX^TM^ *Deluxe Set*, Biolegend^®^, San Diego, CA, USA) according to the manufacturer’s instruction. The samples were read with an absorbance plate reader (Tecan, D300 Digital Dispenser, Männedorf, ZH, Switzerland). The limit of detection was 7.8 pg/mL.

### Primary human cord blood cells

CB-derived cells were isolated from placental tissue and processed by the biobank of the department of Medical Oncology and Hematology, University Hospital Zurich. Following the enrichment of mononucleated cells by density gradient centrifugation, CD34+ cells were purified by CD34 microbeads and LS columns (both Miltenyi Biotec). All primary human cells were cryo-preserved in freezing media. Human cord blood was obtained with parent written informed consent from cords/placentas of healthy full-term newborns (Department of Obstetrics, University Hospital Zurich and Triemli Hospital, Zurich).

### Transgene copy number analysis

Average copy numbers of integrated SB transposons per genome in the polyclonal CAR-T cell populations were determined by droplet digital PCR (ddPCR), as described in ^63^. Briefly, genomic DNA samples prepared from the CAR T cells were subjected to digestion with DpnI to eliminate residual plasmid vector DNA, followed by fragmentation with CviQI. ddPCR reactions were performed by using two primer pairs for the target (the right terminal inverted repeat of the SB transposon) and the reference amplicons, as well as fluorescently labelled hydrolysis probes for both amplicons. Detection of fluorescence and counting of each droplet per ddPCR reaction was done with a droplet reader instrument (BioRad) using a two-color detection system.

### In vivo studies

In vivo studies were conducted with NSG (NOD.Cg-Prkdc^scid^ Il2rg^tm1Wjl^/SzJ) mouse strains. Animal experiments were conducted in compliance with procedures approved by the Veterinäramt des Kantons Zürich, Switzerland (194/2018). The animals were kept in the animal facilities of the University Hospital of Zürich and the Schlieren Campus of Zürich. Generation of humanized mice engrafted with human CB was performed as previously reported ^64^. Briefly, newborn pups were sub-lethally irradiated with 150 cGy with an RS-2000 irradiator (Rad Source, Buford, GA, USA) and transplanted by intrahepatic injection of 1X10^5^ CD34^+^ umbilical CB cells. Levels of chimerism were monitored by flow cytometry of PB post-transplantation. Administration of 2X10^6^ CAR T cells and NT cells was performed by intravenous injections 6-10 weeks post-transplantation. Animals were euthanized 6 days after T cell transfer in the experiment depicted in Figure 1 and 14 days after T cell transfer in the experiment described in Figure 5. For studies with CD117+, luciferase-GFP expressing MOLM-14 cells, adult NSG mice (6-9 weeks of age) were sub-lethally irradiated and intravenously injected with 1X10^5^ MOLM-14 cells. Animals with positive luciferase signal on day 9 were allocated to receive 2X10^6^ CAR+ T cells, corresponding to 7.4X10^6^ SB iC9.CAR T cells, 3.3X10^6^ LV CAR T cells, 7.4X10^6^ NT cells or PBS. For weekly bioluminescent imaging (BLI), mice were anesthetized and received 150 mg/kg bodyweight D-luciferin (PerkinElmer, Inc, Waltham, 148 MA, USA) as intraperitoneal injection. Image acquisition was performed on a Xenogen IVIS 200 machine (Perkin Elmer) with the Living Image® 149 Software. For survival analysis, mice were closely monitored for any sign of suffering and promptly removed from the experiment in case of meeting humane endpoints. Hematological organs (PB, BM, and spleen) were collected and processed for flow cytometry analysis.

### Statistical analysis

Data is represented as mean +/− standard error of the mean. One-way or two-way Anova with Turkey multiple comparison were used to determine the statistical significance of the data. For survival analysis, statistical differences were calculated by Mantel-Cox test. Statistical analyses were performed with GraphPad Prism 5.0 (GraphPad Software, Inc., La Jolla, CA).

## Data availability

Raw data that support the findings of this study that are not present in the manuscript will be made available upon request.

## ACKNOWLEDGMENTS

This work was supported by the Clinical Research Priority Program “Human Hemato-Lymphoid Diseases” of the University of Zurich to MGM; the Swiss Cancer Research (KFS-3846-02-2016) and the Clinical Research Priority Program “ImmunoCure” of the University of Zurich to MGM and DN; the Förderung des Akademischen Nachwuchses (FAN) of the University Hospital Zurich to CFM; a University Research Priority Project Translational Cancer Research grant of the University of Zurich (to RM and SP) and the Swiss National Science Foundation (310030B_166673/1, to MGM). The authors would like to thank Norman Russkamp and Francesco Manfredi for assistance in experiments, Andrea Biondi, and Giuseppe Gaipa for scientific discussion.

## AUTHOR CONTRIBUTIONS

CFM conceptualized the study, designed and perform all experiments, collected and analyzed the data, wrote the manuscript. RM provided assistance in LV preparation and animal experiments with mRNA CAR T cells. SB, MC, MP, LV, CP performed experiments and helped in preparing the figures. SP proposed the use of ivt mRNA for CAR and transposase, designed and produced those mRNA molecules. ZI led the VCN analysis. JS provided assistance in conceptualizing animal experiments. DN conceptualized the study and obtained financial support. MGM conceptualized the study, designed the experiments, obtained financial support, supervised the study, wrote and approved the manuscript. All authors reviewed and approved the final manuscript.

## Supplemental Information

**Supplemental Table 1.** List of fluorescently labeled antibodies/proteins used for flow cytometry.

| Target | Clone | Fluorochrome | Supplier |
| --- | --- | --- | --- |
| Protein-A | n/a | FITC | Thermo Fisher |
| CD3 | OKT3 | APC | BioLegend |
| CD4 | RPA-T4 | FITC | Invitrogen |
| CD8 | SK1 | PerCP-Cy5.5 | BioLegend |
| CD45RA | HI100 | Brilliant Violet 711™ | BioLegend |
| CD62L | DREG-56 | PB | BioLegend |
| CD117 | 104D2 | PE-Cy7 | Thermo Fisher |
| CD27 | M-T271 | BV510 | Biolegend |
| CD95 | DX2 | BV786 | BD Biosciences |
| CD279 | EH12.2H7 | APC Cy7 | Biolegend |
| CD223 | 3DS223H | Alexa-Fluor 700 | BD Biosciences |
| CD366 | F38-2E2 | PE | Ebiosciences |
| mouse CD45 | 30-F11 | PerCP-Cy5.5 | BioLegend |
| CD34 | QBend-10 | PE | Thermo Fisher |
| CD33 | WM53 | Brilliant Violet 711™ | BioLegend |
| CD19 | SJ21C1 | FITC | BioLegend |
| CD45 | HI30 | eFluor™ 450 | Thermo Fisher |
| HLA-A2 | BB7.2 | APC-780 | BD Biosciences |

**Figure S1:**
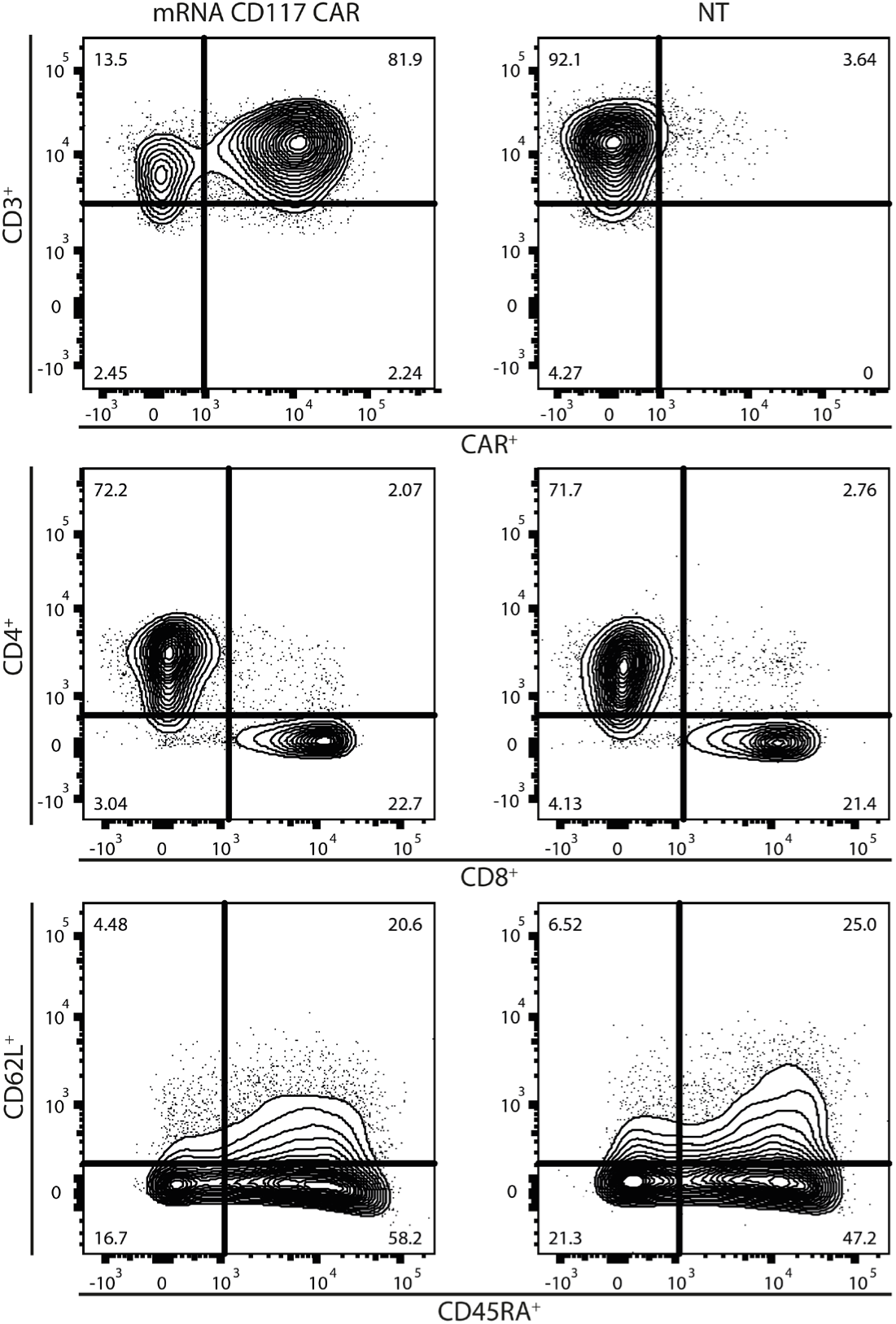
Phenotype of T cells electroporated with mRNA CD117CAR. Purified T cells were stimulated and electroporated with 10 µg of CAR mRNA or in absence of DNA (not transfected (NT)). Flow cytometric immunophenotyping by dual density plots in one representative donor (n = 5). CD3/CAR, CD4/CD8, CD45RA/CD62L expression were measures.

**Figure S2:**
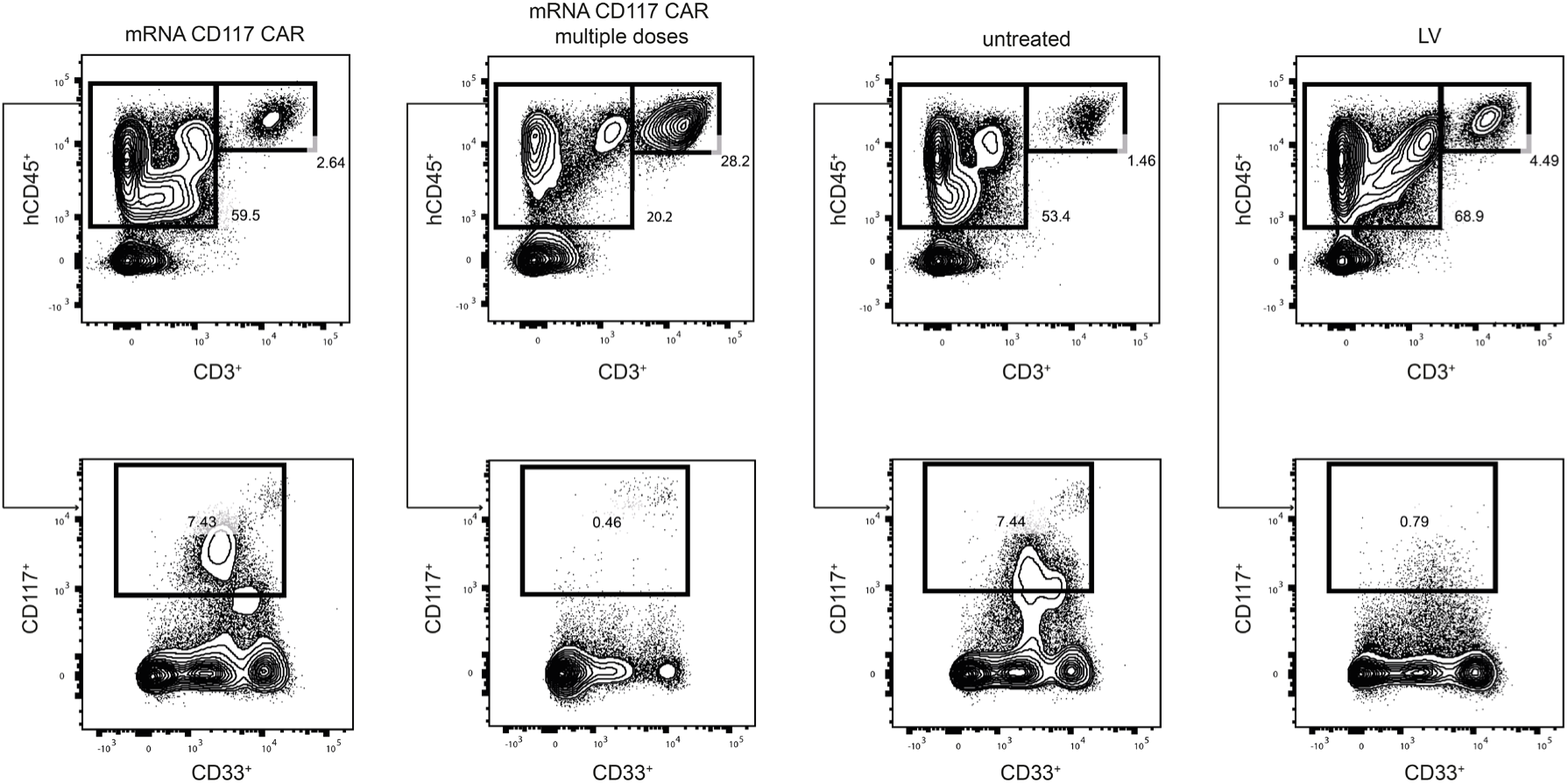
mRNA CD117 CAR T cells deplete healthy CD117 HSPCs in vivo. Newborn NSG mice were sublethally irradiated and injected with CB-derived hCD34+ cells. After having confirmed the engraftment and establishment of human hematopoiesis, mice received a single dose of mRNA CAR T cells (2 x 10^6^), two high doses mRNA CAR T cells (6 x 10^6^ every three days), or a single dose of LV CAR T cells (2 x 10^6^). Representative Flow cytometric immunophenotyping of the bone marrow of treated animals at endpoint. One representative mouse is shown.

**Figure S3:**
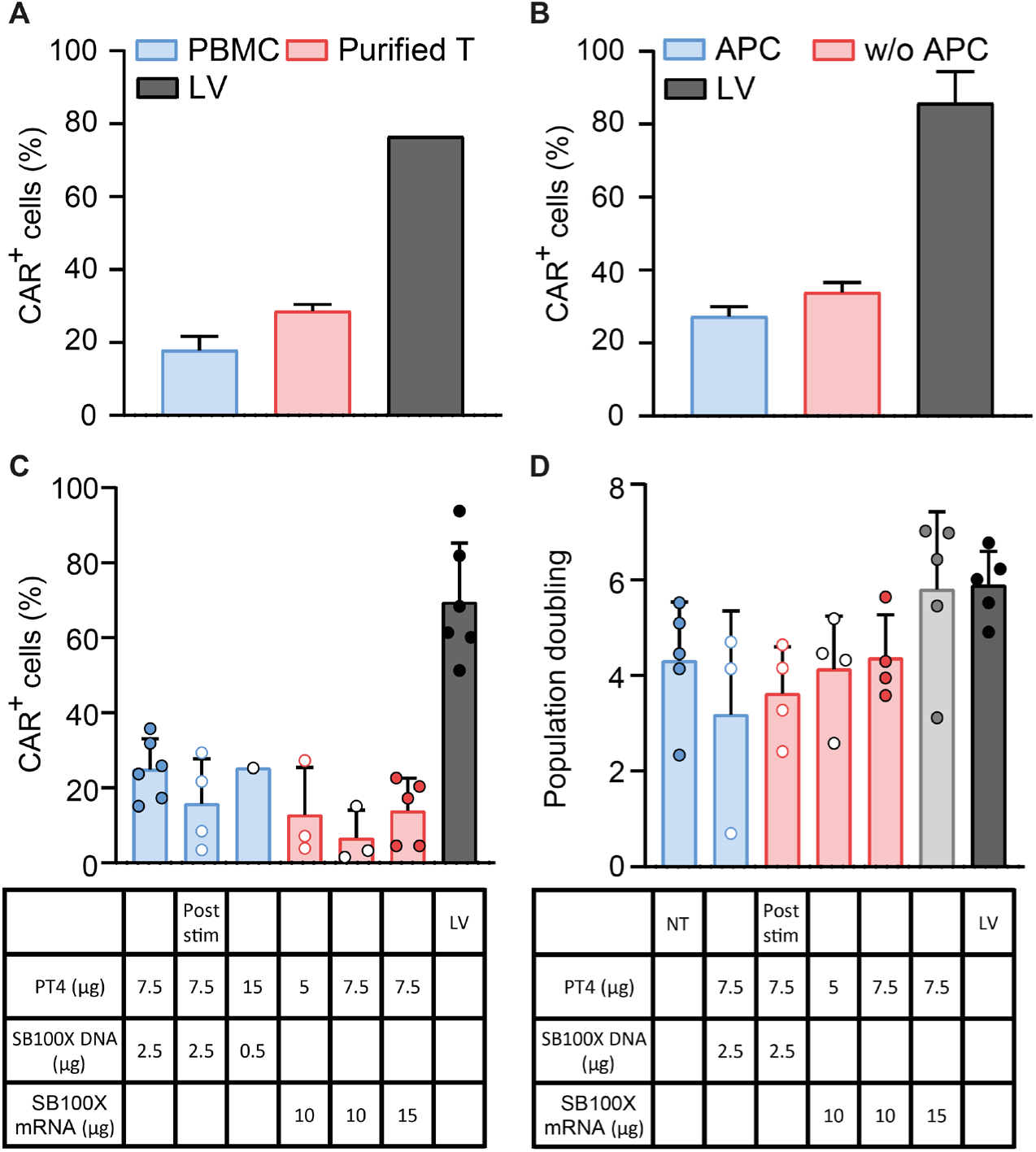
Transduction optimization of CD117 CAR T cells by SB vector. (A) CAR T cells were generated by electroporation of total PBMCs or purified T cells in the presence of SB vector and the SB100X. CAR expression as percentage of CD3 cells as determined by flow cytometry with the recombinant c-Kit protein at 10 days after electroporation. (B) CAR expression as percentage of CD3 cells in T cells electroporated with SB vectors and stimulated in the presence or absence of APC, compared to LV transduced T cells. (C) CAR expression as percentage of CD3 cells of T cells electroporated with different concentrations of the PT4 vector and SB100X transposase, provided as DNA plasmid or mRNA, before or post stimulation, compared to LV transduced T cells, analyzed by flow cytometry at 10 days after electroporation. (D) Population doubling of T cells electroporated with different concentrations of the PT4 vector and SB100X transposase, provided as DNA plasmid or mRNA, before or post stimulation, compared to LV transduced T cells, analyzed by flow cytometry at 10 days after electroporation.

**Figure S4:**
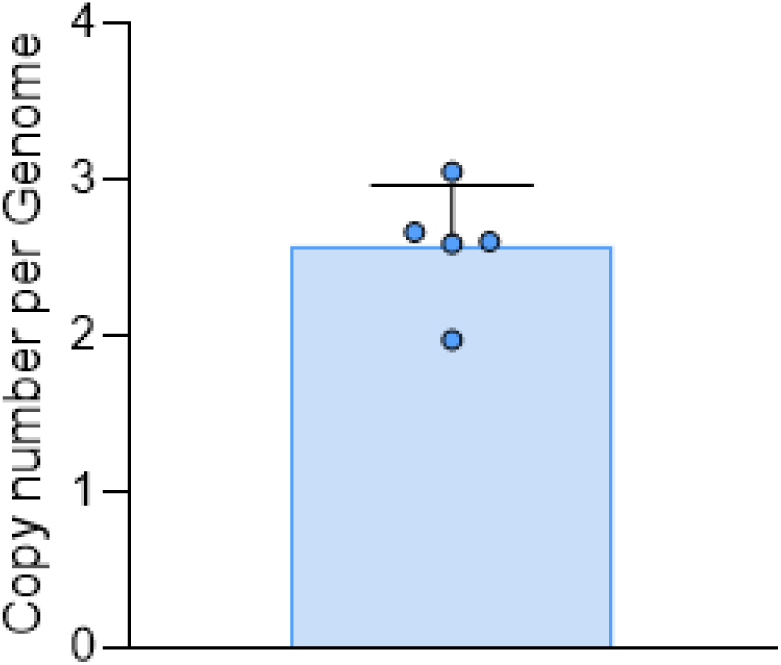
Transgene copy number of CD117 CAR T cells generated by SB vector. CAR T cells were generated by electroporation of total PBMC or purified T cells in the presence of SB vector and the SB100X. Transgene copy number per genome as determined by digital PCR at 10 days after electroporation.

**Figure S5:**
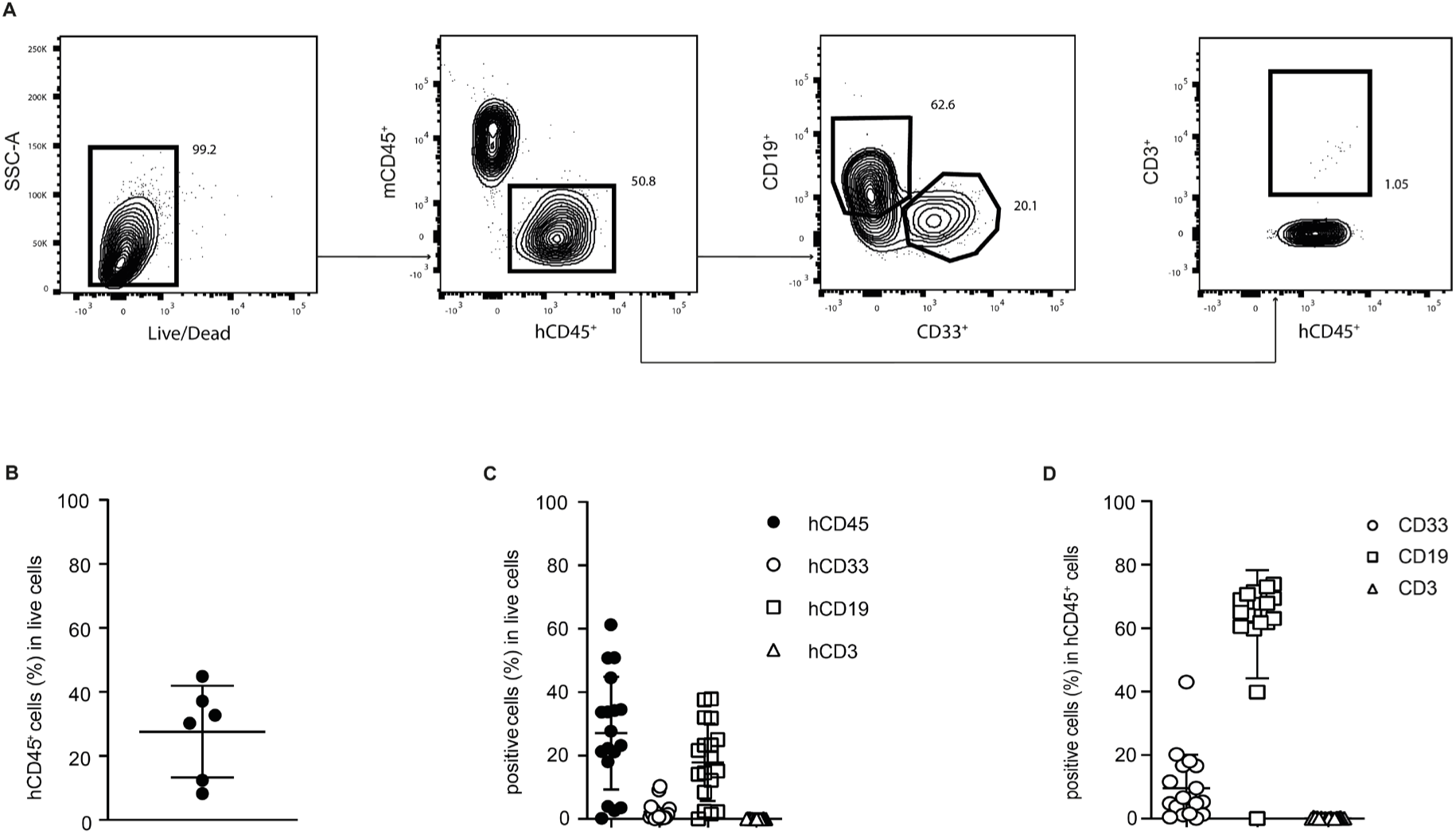
mRNA CD117 CAR T cells deplete healthy CD117 HSPCs in vivo. (A) Representative flow cytometric immunophenotyping with gating strategy of the peripheral blood of mice engrafted with human CD34+ CB cells. One representative mouse is shown. (B) Human engraftment as percentage of hCD45 cells in live cells from peripheral blood of humanized mice, analyzed by flow cytometry at 40 days after transplantation. (C) Human engraftment as percentage of hCD45, hCD33, hCD19, and hCD3 cells in live cells from peripheral blood of humanized mice, analyzed by flow cytometry at 60 days after transplantation. (D) Human engraftment as percentage of hCD33, hCD19, and hCD3 cells in hCD45 cells from peripheral blood of humanized mice, analyzed by flow cytometry at 60 days after transplantation.

